# Effect of prior somatosensory electrical stimulation on the wrist in biasing human hand choice

**DOI:** 10.1101/2023.08.23.554458

**Authors:** Kento Hirayama, Toru Takahashi, Takayuki Koga, Rieko Osu

## Abstract

Hand choice is an unconscious decision frequently made in daily life. A correlation has been found between the state of brain activity before target presentation and hand choice for the target, around which the hand choice probability is in equilibrium. However, whether the neural state before target presentation affects hand choice remains unknown. Therefore, this study aimed to examine whether instantaneous somatosensory electrical stimulation administered to the unilateral wrist at 0, 300, or 600 ms before the target presentation facilitates or inhibits stimulated hand choice for targets around the hand selection equilibrium point. A single electrical stimulation comprised five trains of 1 ms electrical pulses, with a 20 ms inter-pulse interval. The stimulus intensity was set at 80% of the motor threshold. Fourteen right-handed healthy adults (five females, nine males; mean age, 25.1 ± 4.64 years) participated. Unilateral wrist stimulation significantly increased the probability of choosing the stimulated hand and led to a faster reaction time than bilateral wrist stimulation and no-stimulation conditions. The results suggest that prior somatosensory stimulation significantly affects the hand-choice process, effectively promoting selection of the stimulated hand. These findings highlight the potential application of this stimulation method in stroke rehabilitation to facilitate use of the paretic hand.

**Impact Statement:** Prior neural stimulation significantly affected the hand-selection process by promoting the selection of the stimulated hand, thus potentially beneficial in rehabilitating the paretic hand of patients with stroke.

## Introduction

During daily activities, individuals unconsciously select the left or right hand to reach an object, such as a cup. The choice of which hand to use is mainly influenced by target-related information, including the location of the target (which hand can reach it with less effort) and shape and orientation of the target (which hand can grasp it more securely) (Oliveira et al., 2010; Schweighofer et al., 2015; Stoloff et al., 2011). However, the selection rate for each hand reaches equilibrium when the target-related factors are similar for the left and right hands (Schweighofer et al., 2015). The point of subjective equality (PSE) can be estimated as the virtual point in space where participants would have an equal probability of using their right or left hand to reach the target. Reaction time (RT) becomes prolonged around the PSE compared with that outside the PSE, suggesting the presence of competition within the brain (Cisek, 2007; Oliveira et al., 2010). Using single-pulse transcranial magnetic stimulation (TMS), Oliveira et al. (2010) introduced noise to the left posterior parietal cortex (PPC) after the target presentation and before the onset of reaching, the interval when brain competition is likely to occur (Oliveira et al., 2010). A significant reduction in right-hand selection rates and prolongation in RTs were observed for targets positioned around the PSE than for targets not around the PSE, impeding competition resolution. This indicates the potential for intervening in the hand selection process around the PSE. In contrast to the study by Oliveira et al. (2010), the intervention biasing the baseline neural activity at the beginning of competition could potentially facilitate competition resolution and make one hand more likely to be selected over the other, and potentially reduce the competition time. Consequently, the intervened hand may be selected more quickly and with a higher selection probability.

Several studies provide evidence in the field of perceptual decision-making showing that the state of the brain before presentation of a target stimulus, (before the competition starts), correlates with subsequent judgment around PSE (Bode et al., 2012; Hesselmann et al., 2008). Based on the experiment by Bode et al. (2012), participants were required to select between two options: whether the image presented was of a piano or chair. The noise was added to the images, and in one-quarter of the images, there was only noise and no discernable information to determine whether the image was of a piano or chair. Electroencephalogram (EEG) features recorded before image presentation can help to predict the selection outcomes without discriminative information (Bode et al., 2012). Hamel-Thibault et al. (2018) compared the delta waves in the EEG in the motor cortex during target presentation in a task where participants had to select the left or right hand to reach the target. They found that the hand contralateral to the motor cortex, exhibiting a more negative instantaneous delta phase, was more likely to be selected. The negative phase of the delta wave signals an increase in neural excitability, suggesting that hand selection can be predicted based on whether the neural excitability has increased by the time of the target presentation (Hamel-Thibault et al., 2018). They also found a correlation between the RT and delta phase in the motor area. These findings suggest the potential that altering the neural state before target presentation can influence subsequent hand selection; however, intervention studies are required to demonstrate the causal relationship. Hirayama et al. (2021) investigated the effects of 10-min transcranial direct current electrical stimulation (tDCS) on hand selection. They focused on stimulating the left and right PPC. Cathodal stimulation of the left PPC (inhibitory effect) and anodal stimulation of the right PPC (facilitatory effect) were found to increase the probability of left-hand selection around the PSE (Hirayama et al., 2021). Although this suggests that PPC interventions can bias the probability of hand selection, it does not clarify whether the neural state before or after target presentation has an effect. To clarify this point, instantaneous intervention that can change the neural excitability is necessary.

Single pulse TMS to PPC is instantaneous and mainly adds noise and perturbs neural activity as shown by Oliveira et al. (2010); however, peripheral nerve stimulation can induce an instantaneous change in the neural excitability. The electrical stimulation of the median or ulnar nerve at the wrist triggers responses in the primary somatosensory cortex, premotor cortex (PMC), and PPC, with a short latency of approximately 30 ms (Forss et al., 1994; Fujii et al., 1994; Mauguière et al., 1999). The latency for visual information input to the primary visual cortex (V1) is approximately 100 ms. Thus, when somatosensory electrical stimulation is applied to the wrist simultaneously with visual information about the target, the information from the somatosensory electrical stimulation to the wrist is delivered to the cortical area first.

Therefore, this study examined whether short-term somatosensory electrical stimulation applied to the unilateral wrist at the time or immediately before target presentation increases or decreases the probability of selecting the stimulated hand for targets around the PSE, when the selection rate is in equilibrium. Additionally, the study aimed to determine whether it shortens or prolongs the RT during hand selection. The results will help to clarify the causal relationship between neural activity prior to target presentation and hand selection and provide experimental support for the existence of neural competition in decision making. Practically, this work will pave the way for biasing hand selection without conscious or deliberate effort by the individual through a simple and inexpensive method. This could be a step toward stroke rehabilitation to overcome nonuse of the paretic hand in daily life by facilitating paretic hand choice.

## Results

Fourteen right-handed healthy participants were recruited for this study. They were asked to reach the target using the left or right hand as quickly and accurately as possible. The targets were presented at nine random positions on the semicircle (Figure 1).

**Figure 1.**
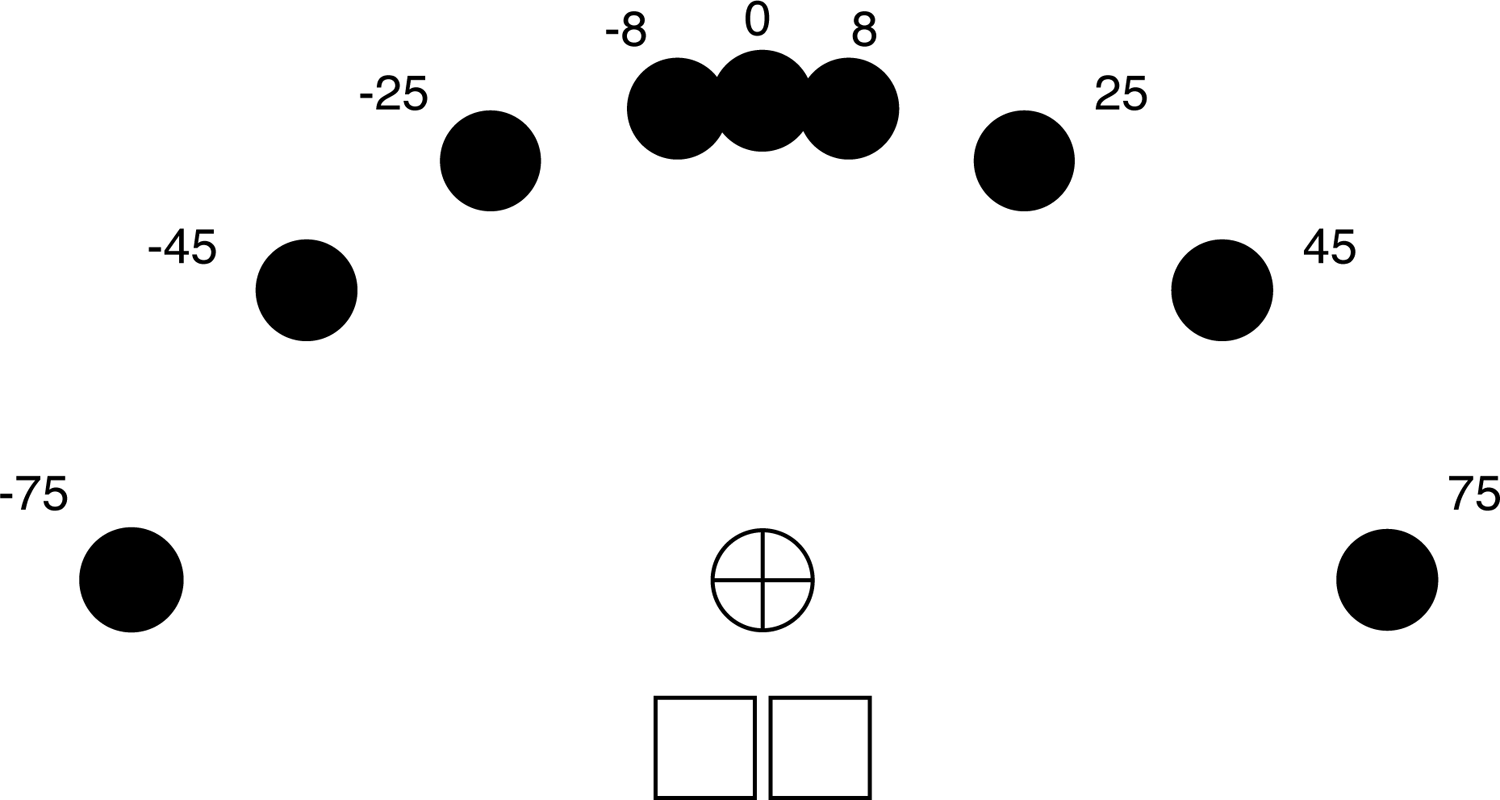
Target positions. Starting position (bottom two squares), fixation point (central “+” mark), and the nine possible targets. The targets were set at angles of 0°, 8°, 25°, 45°, and 75° to either side of the midline of the mirror.

Figure 2 shows that the hand choice probability exhibited a sigmoidal pattern across targets, transitioning from a left-hand preference when dealing with targets on the left side of the space to a right-hand preference when focusing on targets on the right side of the space. The PSE was estimated using logistic regression, representing the point at which participants were equally likely to use their right or left hand to reach targets (Figure 2).

**Figure 2.**
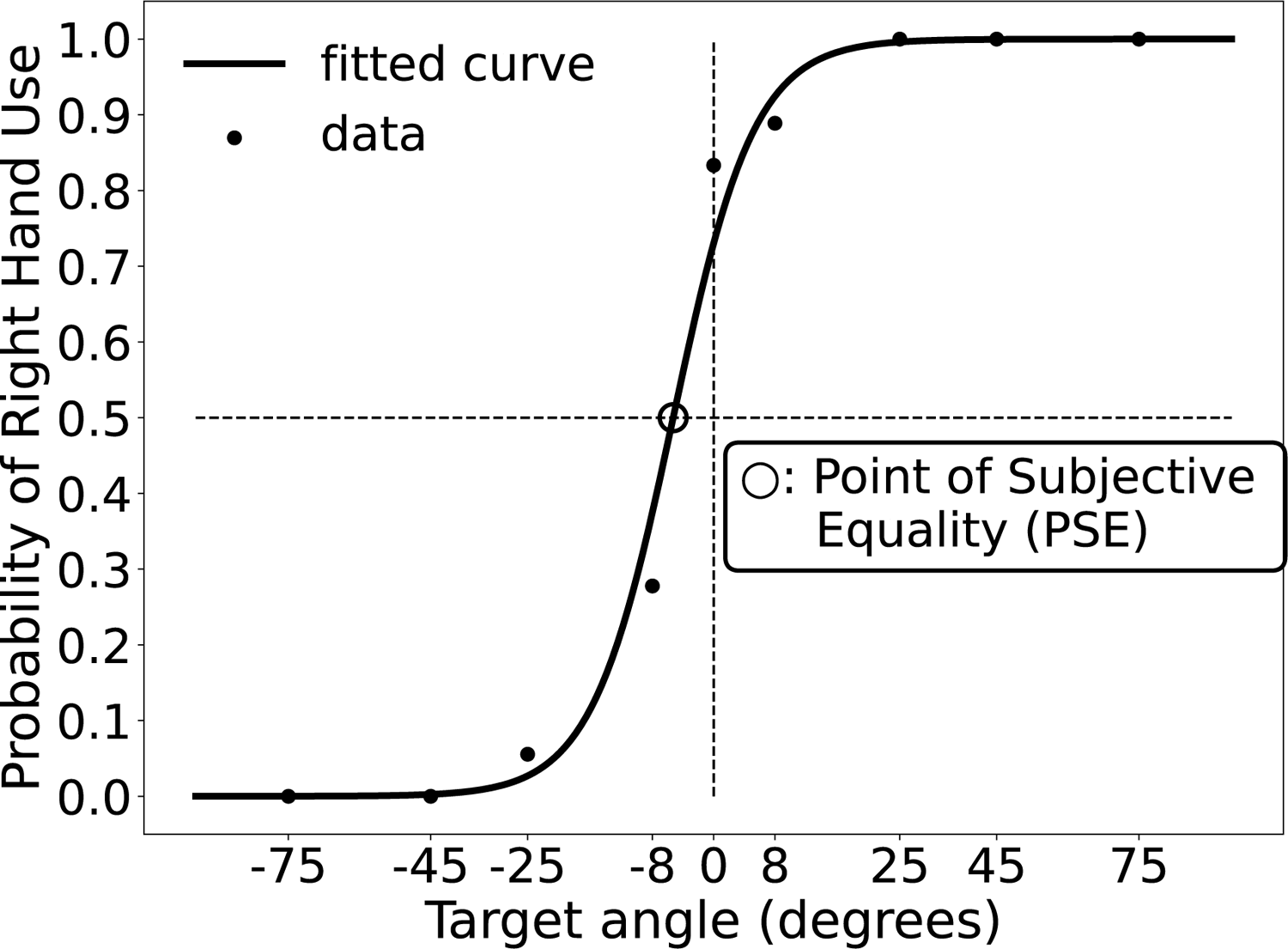
Point of subjective equality (PSE). The probability of choosing the right hand for each target is fitted to a logistic function for each participant. The horizontal axis indicates the target angle, with 0 denoting the midline or physical center (positive degree indicates rightward rotation, negative degree denotes leftward rotation relative to the midline). The vertical axis indicates the rate of right-hand selection. The PSE, indicating the estimated location at which participants were equally likely to use the right and left hands, was thus determined.

The unilateral stimulus (right and left), bilateral stimulus (as the control condition), and no-stimulus conditions were set in this experiment. Stimulation electrodes were attached to the ventral side of each wrist based on peripheral stimulation methods shown to affect sensorimotor regions of the brain in previous studies (Forss et al., 1994; Fujii et al., 1994) (Figure 3).

**Figure 3.**
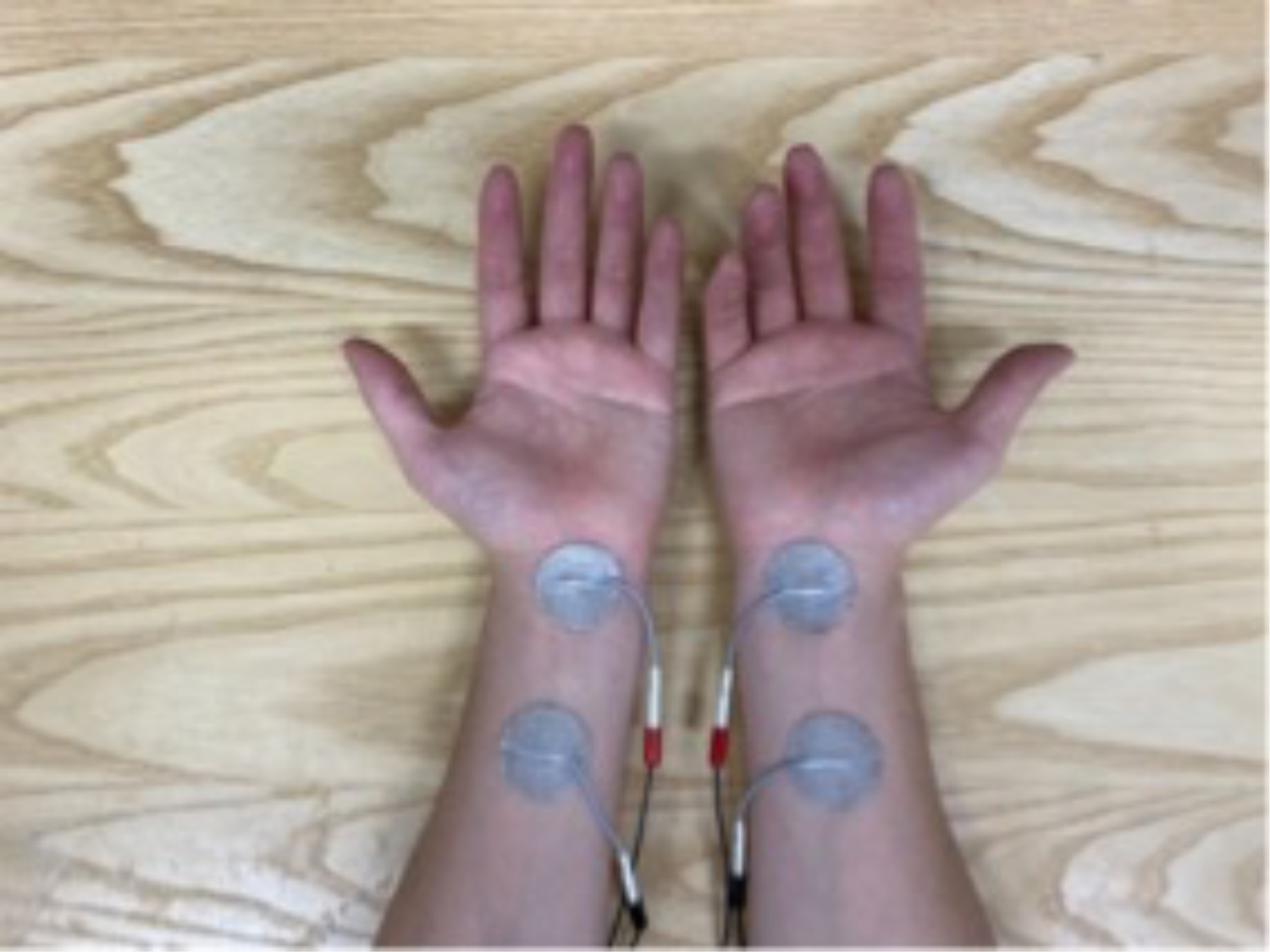
Somatosensory electrical stimulation settings. Anodal electrodes with a 3-cm diameter are attached to the ventral distal end of the forearm on each side, while the cathodal electrodes are positioned 3 cm proximal from the anodal electrodes. The stimulation intensity is 80% of the motor threshold, and the stimulus duration 85 ms. The stimulus conditions are the unilateral stimulus conditions (left and right) and bilateral and no stimulus conditions. Each stimulus condition is set 12 times for each target position and assigned in random order.

The stimulus intensity was set at 80% of the motor threshold, and the stimulus comprised five trains of 1 ms electrical pulses, with a 20 ms inter-pulse interval.

The unimanual reach trials were conducted 648 times, with targets presented 72 times randomly for each position (nine positions). The four stimulus conditions (unilateral [left and right], bilateral, and no-stimulus) were randomly assigned 18 times for each target position. In 36 randomly ordered bimanual reach trials, participants were required to reach for two targets concurrently using both hands. These trials were included to ensure that participants remained prepared to respond with both hands, thereby reducing the likelihood of consistently favoring one over the other. Furthermore, a fixation-reach trial was implemented, where the “+” symbol of the fixation changed to the “×” symbol, and the participants were asked to reach both hands into the fixation circle. There were 36 fixation reach trials in random order to ensure that participants maintained fixation at the start of each trial. The participants were allowed to move their eyes after the target was displayed. The target was presented 0, 300, or 600 ms after the somatosensory electrical stimulation. The intervals between somatosensory electrical stimulation and target presentation were evenly distributed for each target position (nine positions) and stimulus condition (four conditions) (Figure 4).

**Figure 4.**
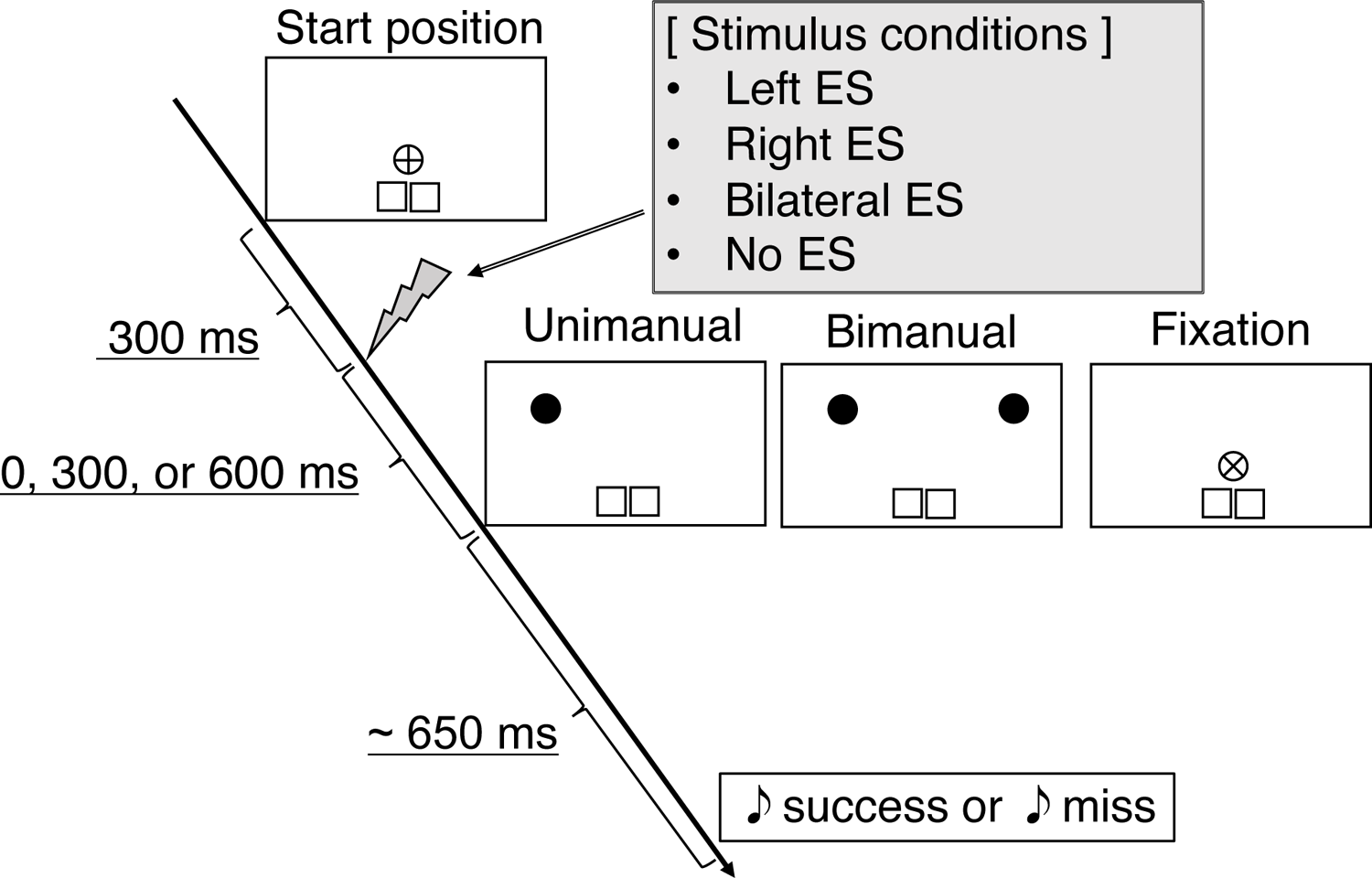
Task protocol. Each trial commenced with each participant pressing two pressure sensors at the starting position with the left and right hands, respectively. After 300 ms, a somatosensory electrical stimulus is administered based on the stimulus conditions. Subsequently, one of three types of tasks is presented randomly at time intervals of 0, 300, and 600 ms after the somatosensory electrical stimulus. In “unimanual reach trials,” the participants reach for the target with one hand. In “bimanual reach trials,” two target circles are presented, and the participant reaches out to them with both hands simultaneously. In “fixation catch trials,” the “+” at the center of the fixation circle is changed to “×,” prompting participants to move both hands to the fixation circle. When one of the sensors on the index finger enters the target circle within 650 ms, the target disappears, and a sound indicating success is played. Otherwise, if the target does not enter the center circle, the target remains visible, and a sound-signaling failure is played.

A logistic function was fitted to the probability of right-hand choice and then averaged across participants for each stimulation condition (Figure 5). In the right (blue line in Figure 5) and left (red line in Figure 5) stimulation conditions, the logistic function exhibited a considerably leftward shift, indicating a preference for the right-hand choice, and rightward shift, indicating a preference for the left-hand choice, respectively, compared with the bimanual and no-stimulation conditions. The effect of the stimulation conditions on the PSE was investigated using a one-way analysis of variance (ANOVA) with repeated measures (Figure 6). A significant main effect was observed in the stimulation condition (*F* (3, 39) = 15.8, *P* < 0.01, partial *η^2^* = 0.55). Multiple comparisons between each pair of stimulation conditions showed that the PSE in the left stimulation condition (PSE = 4.40° ± 10.5°) was significantly larger than that in the other stimulation conditions (*adj P* < 0.05). Conversely, the PSE in the right stimulation condition (PSE = −2.04° ± 9.97°) was significantly smaller than that in the other stimulation conditions (*adj P* < 0.05). There was no significant difference between the PSE in the bilateral stimulation condition (PSE = −0.84° ± 10.6°) and that in the no stimulation condition (PSE = 1.75°±11.3°; *adj P* = 0.16). These findings indicate that the probability of hand choice increased with stimulation. To examine the effect of the timing of the stimuli (0, 300, and 600 ms) before the target presentation on hand selection, the 95% confidence intervals of the PSE for each stimulus timing were calculated using bootstrap with 10,000 resampling of the data to compensate for the limited data (Figure 7). These findings suggest that the closer the stimulus timing was to the target presentation, the probability of choosing the stimulated hand increased.

**Figure 5.**
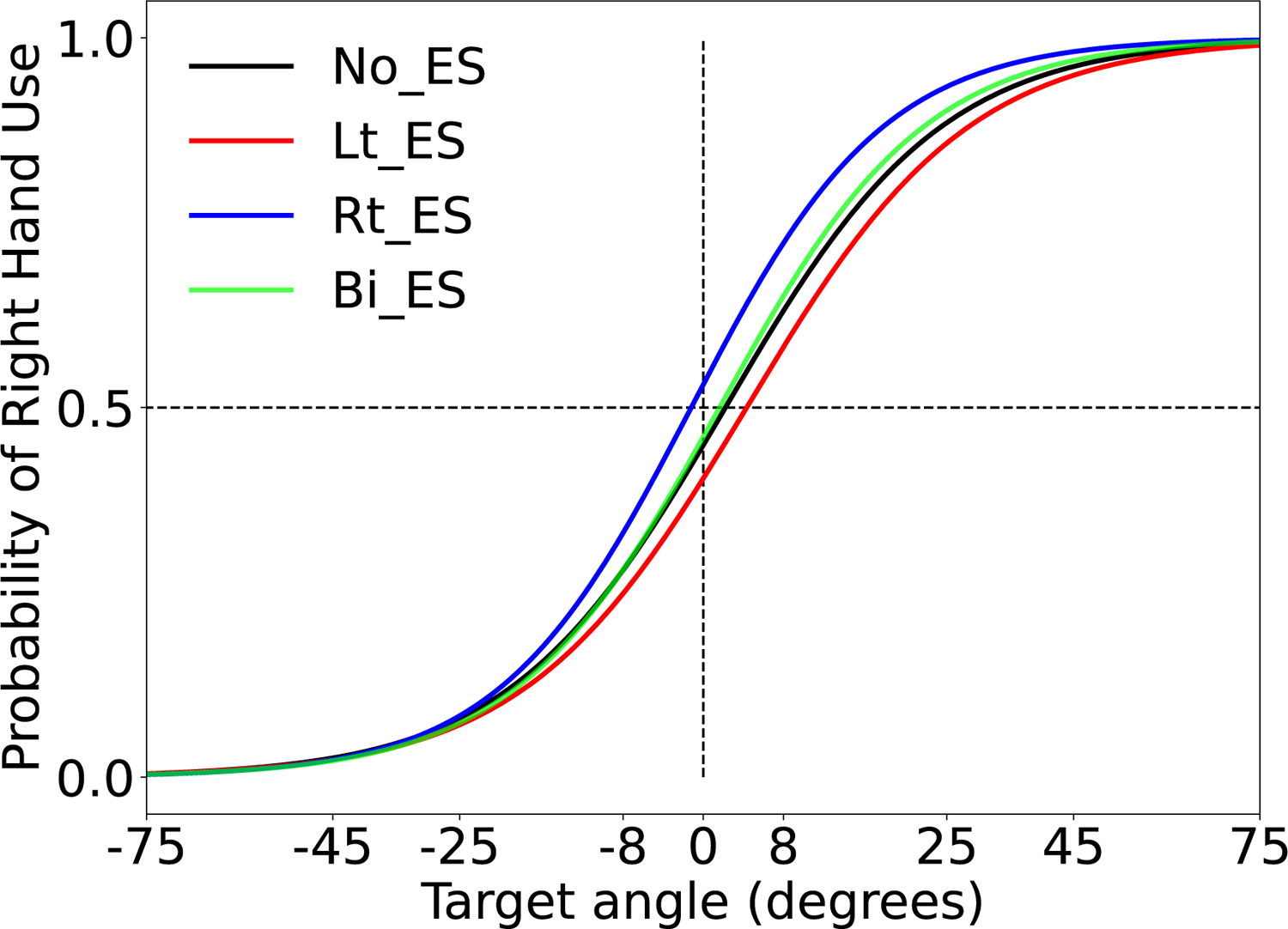
Right hand preference of the participants for each stimulus condition. The logistic function is fitted to the probability of right-hand choice and then averaged across the participants for each stimulus condition. The vertical axis indicates the probability of the right hand-choice, while the horizontal axis represents the target presentation angle. The angle of 0° corresponds to the body-centered angle. Each curve depicts an increase in left-hand choice as the angle shifts to the right side and an increase in right-hand choice as the angle shifts to the left side. The black line represents the no stimulus condition (No_ES), the red line indicates the left stimulus condition (Lt_ES), the blue line indicates the right stimulus condition (Rt_ES), and the green line represents the bilateral stimulus condition (Bi_ES).

**Figure 6.**
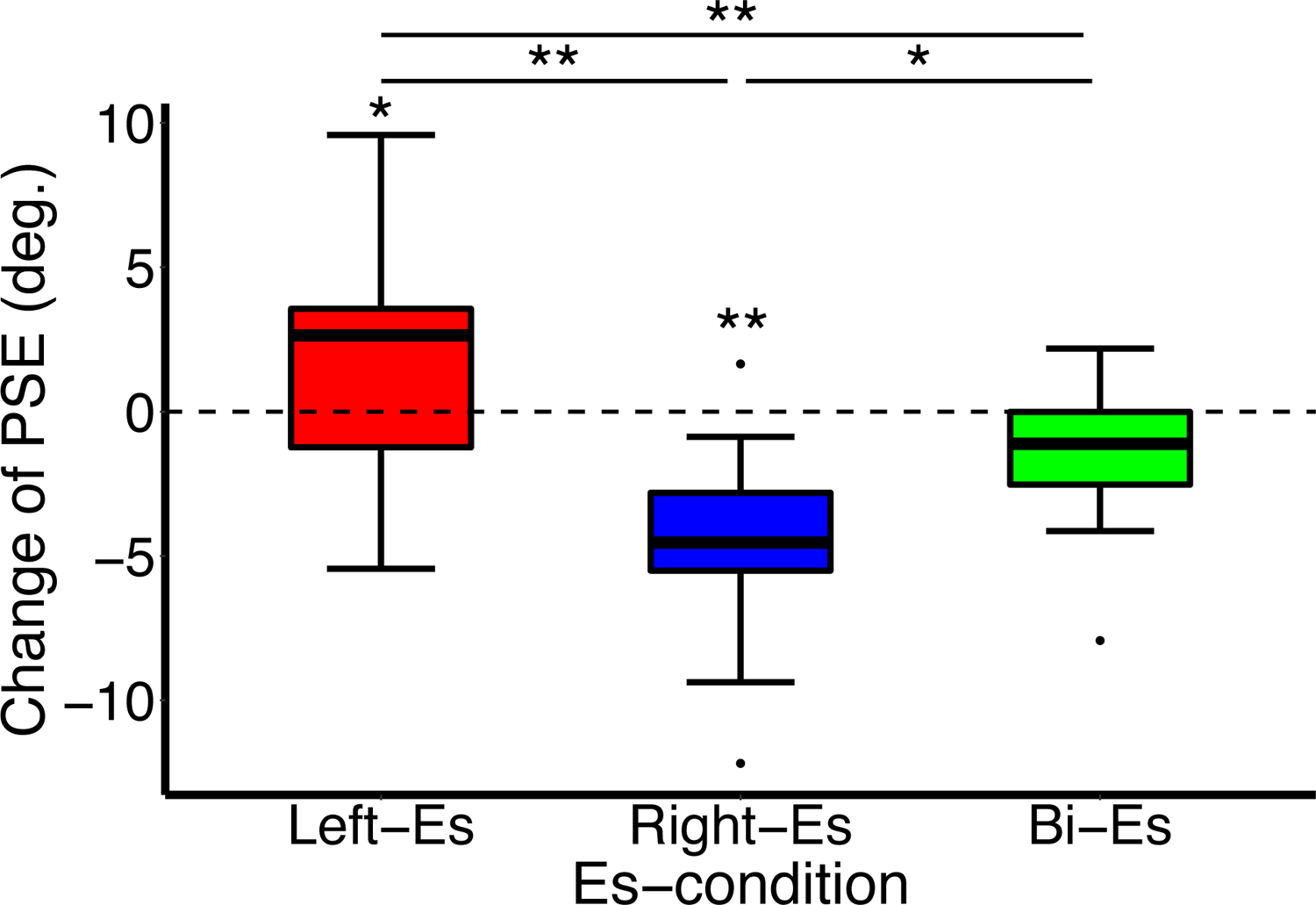
Change in the point of subjective equality (PSE) for each stimulation condition from the no-stimulus condition. The red (Left-Es), blue (Right-Es), and green boxes (Bi-Es) represent the left, right, and bilateral stimulus conditions, respectively. The vertical axis indicates the PSE; 0° represents the PSE of the no-stimulus condition. Larger angles indicate increased left-hand choice, while smaller angles depict increased right-hand choice. The horizontal axis indicates the stimulation condition (Es-condition). **: *P* < 0.01, *: *P* < 0.05.

**Figure 7.**
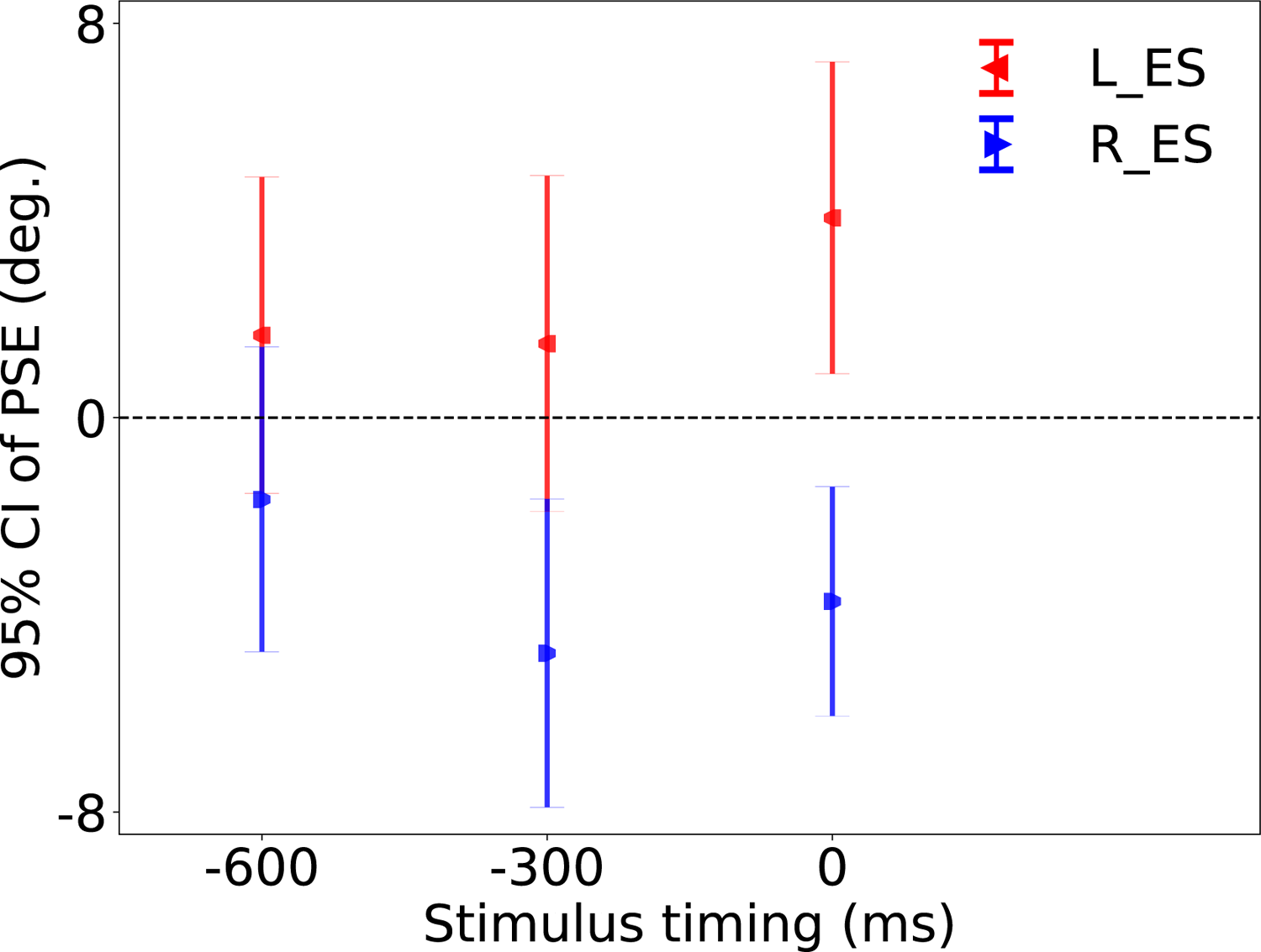
PSE 95% confidence interval for each stimulus timing in the left and right stimulation conditions. The red left-oriented triangles indicate the left stimulus condition, while the blue right-oriented triangles represent the right stimulus condition. The vertical axis indicates the 95% confidence interval of the PSE for each condition, with 0° representing the angle of the body center. A larger value indicates an increase in left-hand choice and a smaller value depicts an increase in right-hand choice. The horizontal axis indicates stimulation timing based on the target presentation time.

To examine whether somatosensory electrical stimulation facilitated or inhibited competition resolution for hand choice, the median RTs were analyzed for five center targets (ranging from −25° to +25°), where hand selection is in equilibrium, and for the other extreme targets (± 75°, ± 45°). These RT values were then compared across each stimulation condition. The RT was defined as the duration between the target presentation and release of the selected hand from the pressure sensor. A one-way ANOVA with repeated measures was conducted to examine the influence of the simulation condition on the center and extreme target RTs. Regarding the center RT results (Figure 8A), a significant main effect was observed in the stimulation condition (*F* (2.11, 27.4) = 3.48, *P* < 0.05, partial *η^2^* = 0.21). In the multiple comparisons between each pair of stimulation conditions, the RT in each of the left (RT = 431.9 ± 25.4 ms) and right stimulation conditions (RT = 429.8 ± 27.4 ms) was significantly smaller than that in the no stimulation condition (RT = 437.1 ± 28.5 ms) (*adj P* < 0.05). However, the RT of the bilateral stimulation condition (RT = 434.2 ± 28.8 ms) was not significantly different from the no stimulation condition (*adj P* = 0.60). In addition, the RT was not significantly different between the left and right stimulation conditions (*adj P* = 0.61). Regarding the extreme target RT results (Figure 8B), there was no significant main effect observed in the stimulation condition (*F* (2.18, 28.35) = 0.91, *P* = 0.42, partial *η^2^* = 0.06). The RT for no, left, right, and bilateral stimulation conditions were 428.1 ± 20.9 ms, 426.4 ± 19.7 ms, 424.2 ± 23.1 ms, and 427.3 ± 20.4 ms, respectively.

**Figure 8.**
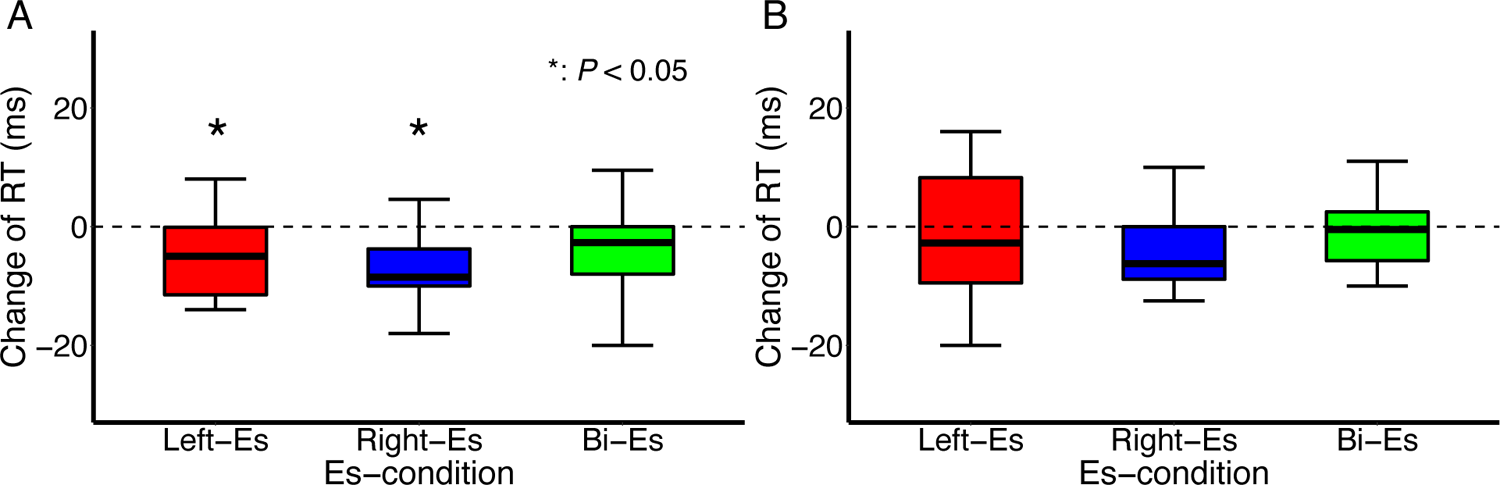
Changes in the reaction time (RT) of the center target (A) and extreme target (B) relative to the no stimulus condition in each stimulation condition. The red (Left-Es), blue (Right-Es), and green boxes (Bi-Es) represent the left, right, and the bilateral stimulus conditions, respectively. The vertical axis represents the change in the RT (ms), while the horizontal axis indicates stimulus conditions (Es-condition). The horizontal line inside each box represents the median. The boxes extend to the lower and upper quartiles, whereas the whiskers extend to the extreme values other than outliers. *: *P* < 0.05.

## Discussion

Somatosensory electrical stimulation of the left or right unilateral wrist at or immediately before the target presentation increased the probability of choosing the stimulated hand. Furthermore, when the target was positioned around the center, the RT was significantly shorter in the unilateral wrist than in the bilateral wrist and no-stimulation conditions.

Somatosensory electrical stimulation was administered in this study at times of 0, 300, or 600 ms before the target presentation. Electrical stimulation to the wrist can influence the neural processes involved in hand selection before the target visual information reaches the brain, considering the latency of the visual-evoked potentials. Furthermore, the transition of somatosensory information from peripheral tissues follows a pathway that reaches the primary somatosensory cortex before projecting to the PPC and PMC. Forss et al. (1994) used magnetoencephalography to measure somatosensory-evoked potentials (SEP) resulting from electrical stimulation of the median nerve at the wrist, specifically at a motor threshold intensity (Forss et al., 1994). They found that an early component of the SEP appeared in the primary somatosensory cortex and PPC with a latency of approximately 20–30 ms. Similarly, Fujii et al. (1994) used electroencephalography to investigate the effect of electrical stimulation of the median nerve at the wrist. They observed the SEP in the contralateral PPC (P3,4) and PMC (F3,4), with a latency of approximately 30 ms (Fujii et al., 1994). The stimuli in this study produced the SEP peaking at approximately 30 ms in the C4 region, suggesting that the stimuli also reached the PPC and PMC. In contrast, the visually presented target information is processed through a different pathway. It reaches the V1 and transferring to the PPC and PMC. Visual information usually requires approximately 100 ms to reach the V1 (Walsh et al., 2005). Evoked potentials are observed around the frontal/parietal regions with a latency of approximately 150–200 ms after the onset of the visual stimulus (N155/N180) (Beck et al., 1980; Di Russo et al., 2002). These previous studies suggest that the somatosensory stimulation reached the PPC and PMC before the target information arrived. Therefore, sensory stimuli seem to have influenced the brain state before the hand-selection process was triggered by the target information, even in cases where sensory stimuli were presented simultaneously with the target.

Researchers have suggested that hand choice exhibits stochastic behavior owing to competing neural activities arising from similar reaching costs. There are two theories of neural competition: one assumes that competition occurs in the PMC (Cisek, 2007; Hamel-Thibault et al., 2018) and the other in the PPC (Fitzpatrick et al., 2019; Oliveira et al., 2010). The somatosensory stimulation in this study is believed to have likely reached both the PMC and PPC, which may have influenced neural competition in either the PMC, PPC, or both regions. Neural competition in the PMC was studied by Hamel-Thibault et al. (Hamel-Thibault et al., 2018). They examined the delta wave in the EEG during target presentation in a task that required participants to choose between their left or right hand to reach the target. They reported that the hand contralateral to the motor cortex, exhibited a more negative instantaneous delta phase and was more likely to be selected. Since the negative phase of the delta wave indicates an increase in neural excitability, hand selection can be predicted by whether the PMC excitability had increased by the time of the target presentation (Hamel-Thibault et al., 2018).

Conversely, there was a functional magnetic resonance imaging (fMRI) study on neural competition in the PPC by Fitzpatrick et al. (Fitzpatrick et al., 2019). They measured brain activity during a hand-choice task. They reported increased activity in the contralateral PPC of the selected hand in the choice condition compared with the instructed condition, where participants reached the target with a pre-instructed hand. Furthermore, Oliveira et al. (2010) provided a momentary noise stimulus to stimulate PPC activity during hand selection by single-pulse TMS 100 ms after target presentation (Oliveira et al., 2010). They reported that TMS of the left PPC significantly reduced the probability of choosing the right hand. Based on the findings of previous studies, there is a possibility that the somatosensory stimulation in this study influenced the neural competition in either the PMC, PPC, or both, thereby facilitating the selection of the stimulated hand. However, it is unclear whether the PMC or PPC is primarily affected by somatosensory stimulation since it reaches both areas. Further studies are required to confirm this hypothesis.

The RTs for targets near the center were prolonged compared with those for targets at the edges in all stimulus conditions in this study. These results are consistent with that of Oliveira et al. (2010), who demonstrated that the RTs to targets located near the center, where hand choice is in equilibrium, are prolonged compared with those for targets at the edges (Oliveira et al., 2010). Those results suggest that the selection competition is larger for hand selection on targets near the center than that for those at the edges. RT is often used to indicate difficulty in the decision-making process. Previous studies have reported that the more equilibrium of the determinants between alternatives, the longer the RT in tasks involving the choice of one option from multiple alternatives (Christopoulos & Schrater, 2015; Julie et al., 2010; Ratcliff & McKoon, 2008). In this study, RTs near the center were significantly shorter in the unilateral wrist stimulation condition than in the no-or bimanual stimulation conditions. However, no significant differences in the RTs were observed among the four stimulus conditions for edge targets. Therefore, stimulation of the unilateral wrist facilitated decision making only when conflicts among alternatives were large, not when conflicts were small because determinants or costs were sufficiently different.

The phenomenon of prolonged RTs in situations in which the choice is in equilibrium has been explained using decision-making models. The Drift Diffusion Model of perceptual decision-making describes the neural activity related to each alternative and explains that the alternative, whose neural activity reaches the choice threshold, is selected first (Ratcliff & McKoon, 2008). The rate of increase in this neural activity is slower than in the competitive state, thus prolonging the time required to reach the threshold and increasing the RT. In action selection involving movement, neural activity occurs for each of the alternatives and compete with each other (Cisek & Kalaska, 2005; Cui & Andersen, 2007); this was demonstrated using the Affordance Competition Model (Cisek, 2007). According to these decision-making models, the factors influencing choice and the RT include the base state of neural activity before the presentation of the target stimulus and the rate at which neural activity increases. Oliveira et al. (2010) stated that interrupting this process by intervening in hand selection with single-pulse TMS to the PPC after target presentation leads to longer RTs (Oliveira et al., 2010). However, the electrical stimulation in this study, applied before or at the target presentation, shortened the RT, suggesting that neural activity reached the threshold of choice earlier, possibly owing to the bias in the base state before target stimulus presentation or the impact on the rate of increase after target stimulus presentation, or both.

In this study, the application of electrical stimulation just before the target presentation had a biased effect on neural activity, reducing competition and facilitating the subsequent hand choice. However, in real-world applications, it is unknown when a target appears. In such a scenario, a potential solution could involve consistent bias in the neural activity that affects hand choice, regardless of the timing of the target presentation. Previous studies have reported that repetitive somatosensory electrical stimulation on the wrist, lasting for 1.5 to 2 hours, resulted in a sustained increase in neuronal activity in the sensorimotor area (Kaelin-Lang et al., 2002; Ridding et al., 2001; Wu et al., 2005). Wu et al. (2005) performed electrical stimulation with a motor subthreshold intensity of the median nerve at the wrist for 2 hours in healthy participants and examined the sensory-motor area activity using fMRI (Wu et al., 2005). They reported a significant increase in primary sensory and motor area activity for up to 60 minutes after median nerve stimulation compared with the condition without stimulation. However, whether the repetitive wrist stimulation affects the PPC or PMC has not been investigated.

Non-invasive brain stimulation could be another option that consistently biases the state of neural activity related to hand selection. Previous studies using tDCS on the PPC or primary motor area have revealed a residual effect on hand selection (Hirayama et al., 2021; Javadi et al., 2015). Hirayama et al. (2021) used tDCS to continuously change the PPC activity during a hand-selection task (Hirayama et al., 2021). The results showed a significant increase in left-hand selection after stimulation compared with pre-stimulation when placing the cathodal electrode on the left PPC and the anodal electrode on the right. In addition, Javadi et al. (2015) performed a perceptual decision-making task where participants were asked to look at a rectangular figure and to respond with their left hand when the vertical side was longer and with their right hand when the horizontal side was longer (Javadi et al., 2015). During the task, tDCS simultaneously promoted or inhibited activity in the left and right primary motor areas. They reported that when the stimulus was a square, the hand contralateral to the primary motor area, whose activity was facilitated, was more likely to be selected. Hirayama (2021) and Javadi (2015) did not clarify the effective time point of the hand selection process; however, the effect may potentially be due to changes in the base brain state, which was the focus of this study.

This study had a limitation. The specific brain regions where neural activity was altered by the stimulus and influenced hand selection were unclear. It is necessary to examine the relationship between changes in neural activity and hand selection using neurophysiological techniques, such as EEG, or neuroimaging techniques, such as fMRI.

The present findings hold potential applications in rehabilitation, as they may contribute to promoting the use of the hemiparetic hand in patients with stroke who experience reduced quality of life. Patients with stroke may regain the ability to use their paretic hand through rehabilitation; however, they often rely on their nonparetic hand for daily activities, resulting in “learned nonuse,” indicating use of the paretic hand is significantly reduced. (Han et al., 2013; Taub et al., 2006). These findings suggest that somatosensory electrical stimulation of the unilateral hand before target presentation can unconsciously prompt a stimulated hand choice. A possible rehabilitation application would be to apply stimulation to the paretic hand and increase the probability of paretic hand choice in the daily life of the patient. Future studies should examine the effectiveness of the proposed method in promoting the use of the paretic hand of patients with stroke.

## Materials and Methods

### Participants

Overall, 14 right-handed healthy participants (five females, nine males; mean age, 25.1 ± 4.64 years) were recruited in this study; they provided written informed consent and were remunerated for their participation. The required sample size was calculated using G*Power v3.1, with a power of 0.9, an alpha level of 0.05, and a large effect size of 0.4. Following the power analysis, a sample size of 13 was required for a one-way repeated-measures ANOVA with four measurements. The inclusion criteria were as follows: (1) no history of nerve or orthopedic injuries to the upper extremity and (2) no history of chronic or acute neurological, psychiatric, or medical illnesses. The study was approved by the Waseda University Ethics Committee (Tokyo, Japan) and performed in accordance with the principles of the Declaration of Helsinki.

### Experimental setup

The participants were comfortably seated on chairs, with their hands resting on the tabletop surface. A horizontal display was placed above the table, and the mirror was placed midway between the display and table surface. The height of the mirror was set in a way that allowed the participants to see their reflection and not the display or their hands. The stimulus was presented on the display, and its reflection in the mirror gave the impression that it was on the table. To track the motion, two or three-dimensional motion tracking sensors (Fastrak, Polhemus, Colchester, VT, USA) were attached to the index finger of each participant, and the fingertip positions were measured at a sampling rate of 60 Hz. In addition, the current position of both hands was indicated through feedback in the form of two black dots. The participants were instructed to position their fingertips on the pressure sensors placed on the table. This setup allowed the sensors to detect when the hands of the participants were released from the table. A circle with a “+” symbol was displayed 2.5 cm away from the center to inform the participants of the target visual fixation position (Figure 1).

### Task protocol

At the beginning of each experiment, the participants were instructed to press the pressure sensor and to focus their eyes on the fixation point (Figure 4). After 300 ms, somatosensory electrical stimulation was applied to the unilateral wrist (left or right), bilateral wrist, or no stimulation at all, in random order (detailed settings for the somatosensory electrical stimulation are described below). One of three different trials was initiated at random intervals of 0, 300, or 600 ms after the somatosensory electrical stimulation. In the “unimanual reach trial,” a target circle with a diameter of 4 cm would appear at one of nine positions on a semicircle approximately 27 cm away from the starting position. The targets were presented randomly at these nine positions. The participants were instructed to reach the target as quickly as possible with either hand within 650 ms after it appeared. The hand that was not selected needed to be held at the starting position. A disappearing target and sound indicated success if the index finger sensor was within the radius of the target in 650 ms. However, if the index finger sensor failed to enter the radius of the target within 650 ms, a sound indicating failure was played as the target remained visible. The “bimanual reach trial” required participants to simultaneously reach two targets on a semicircle with both hands within 650 ms to reduce the probability of using the same hand for any target. The “fixation reach trial” involved changing the “+” mark to an “x” mark, where participants had to place the index fingers of both hands in the fixation circle within 650 ms to ensure fixation before the target presentation. Participants were allowed to move their gaze freely after target presentation and were instructed to return their hands to the starting positions after feedback sounds. No electrical stimulation was administered during the “bimanual reach” and “fixation reach trials.”

### Procedure

One block consisted of 120 trials, including 108 unimanual (nine targets on the semicircle displayed 12 times), six bimanual, and six fixation reach trials. These were presented in a pseudo-random order, and the participants performed six blocks. Before starting the task, participants completed three practice sessions, each consisting of 20 trials. The practice sessions consisted of two predetermined hand-choice sessions (only the right or left hand) in random order, followed by a free-choice session.

### Somatosensory electrical stimulation settings

We used circular electrodes with a diameter of 3 cm along with an electrical stimulator (Nihon Koden Corporation, Tokyo, Japan). The anodal electrodes were attached to the ventral distal end of the forearm on each side, whereas the cathodal electrodes were positioned 3 cm proximal to the anodal electrode (Maharjan et al., 2018). Stimulation intensity was set at 80% of the motor threshold, which was determined as the minimum intensity required for a visible thumb twitch. Each somatosensory electrical stimulation consisted of five consecutive 1-ms rectangular stimuli with an interval of 20 ms (50 Hz) (Figure 9).

**Figure 9.**
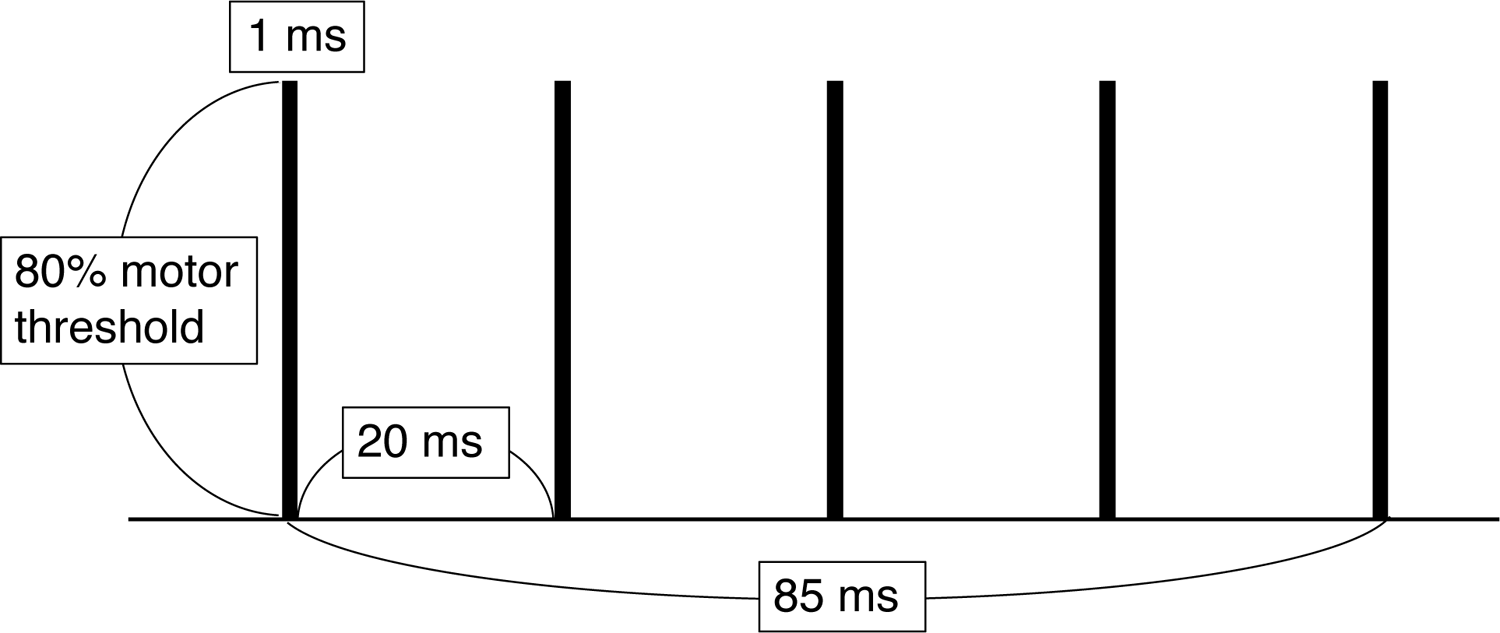
Details of a somatosensory electrical stimulus. A somatosensory electrical stimulation consisted of five consecutive 1-ms rectangular stimuli with an interval of 20 ms. The total stimulus duration is 85 ms. The stimulus intensity is set at 80% of the motor threshold.

### Analyses and statistics

#### Hand choice probability

The hand that had reached a distance of 2.5 cm away from the starting position before the other hand was considered the selected hand. Hand preference was evaluated by calculating the probability of the right choice in each stimulus condition, where the right choice was assigned one, and the left hand was assigned zero for each position that presented the target. Using a logistic regression model, we determined that the angle of the target position, resulting in a 50% right-hand choice probability, represented the PSE (Figure 2).

Furthermore, to investigate the effect of the stimulation condition on the PSE, we performed a one-way repeated measures ANOVA. In case of a statistically significant main effect, we performed multiple comparisons between stimulus conditions using a paired t-test corrected by Shaffer. In addition, a bootstrap method was employed with 10,000 resampled data points from all participants to calculate the 95% confidence interval of the PSE for each stimulus timing (0, 300, and 600 ms). This was conducted to compensate for the relatively small amount of data available (84 and six data points for each participant for each target position).

#### Reaction time

The RT was defined as the duration between target presentation and the release of the selected hand from the pressure sensor. The median RT was determined for five center targets ranging from −25° to +25° (center target RT) and four extreme targets positioned at ± 45° and ± 75° (extreme target RT). A one-way repeated measures ANOVA was used to examine the effect of the stimulation condition on the center and extreme target RTs. When a statistically significant main effect was observed, multiple comparisons between stimulus conditions were performed using a paired t-test corrected by Shaffer. Furthermore, to begin each statistical analysis, the Shapiro–Wilk test was performed to confirm data normality. Mauchly’s sphericity test was used to test the spherical assumption, and if it was rejected, the degrees of freedom were adjusted using the Greenhouse-Geisser method. Statistical analyses were performed using the R software (version 3.6.1; The R Foundation, Vienna, Austria). A statistical significance level was set at *P* < 0.05.

## Acknowledgments

The authors are grateful to Mr. Yuta Takahashi, a member of the Cognitive Neuroscience Lab, for helpful discussions.

## Data Availability Statement

The datasets generated and analyzed during the current study are available from the corresponding author on reasonable request.

## Author details

Kento Hirayama, Faculty of Human Sciences, Waseda University, Saitama, Japan Contributions: Conceptualization, Methodology, Formal analysis, Investigation, Writing—original draft, writing —review and editing, funding acquisition, Toru Takahashi, Faculty of Human Sciences, Waseda University, Saitama, Japan Contribution: Methodology, Investigation, Writing—review & editing, Takayuki Koga, Graduate School of Human Sciences, Waseda University, Saitama, Japan Contribution: Formal analysis, Rieko Osu, Faculty of Human Sciences, Waseda University, Saitama, Japan Contribution: Conceptualization, Methodology, Supervision, Writing—review and editing, and funding acquisition For correspondence

